# Development of an *in-situ* printing system with human platelet lysate-based bio-adhesive to treat corneal perforation

**DOI:** 10.1101/2021.09.13.460167

**Authors:** Jingjing You, Hannah Frazer, Sepidar Sayyar, Zhi Chen, Xiao Liu, Adam Taylor, Benjamin Filippi, Stephen Beirne, Innes Wise, Chris Hodge, Gordon Wallace, Gerard Sutton

## Abstract

**Purpose:** Corneal perforation is a clinical emergency. Tissue glue to seal the perforation, and supplementary topical medication represents existing standard treatment. Previously, our group developed a transparent human platelet lysate (hPL)-based biomaterial that showed good cell compatibility and accelerated corneal epithelial cells healing *in-vitro.* This study aims to develop a novel treatment method for corneal perforation using this biomaterial.

**Methods:** Rheometry was used to measure the hPL-based biomaterial behaviour at room and corneal surface temperatures. Its adhesiveness to porcine skin and burst pressure limit were also measured. Based on rheological behaviour, a hand-held biopen was developed to extrude it onto the cornea. An animal trial (5 New Zealand white rabbits) to compare impact of the biomaterial and cyanoacrylate glue (control group) on a 2mm perforation was conducted to evaluate safety and efficacy.

**Results:** The hPL-based biomaterial showed higher adhesiveness compared to commercial fibrin glue and withstood burst pressure approximately 6.4× higher than routine intraocular pressure. Treatment rabbits had lower pain scores and faster recovery, despite generating similar scar-forming structure compared to controls. No secondary corneal ulcer was generated in rabbits treated with the bio-adhesive.

**Conclusions:** This study reports a novel *in-situ* printing system capable of delivering a hPL-based, transparent bio-adhesive and successfully treating small corneal perforations. Bio-adhesive-treated rabbits recovered faster and required no additional analgesia. Both groups showed scarred corneal tissue after healing, however no infection and inflammation was observed by 3 weeks. The delivery system was easy to use and may represent an alternative treatment for corneal perforation.

**Highlights:**

- This study presents a novel *in situ* printing system to treat corneal perforation
- The system is comprised of a human platelet lysate-based bio-adhesive and a pen-like hand held delivery system
- Mechanical tests showed our transparent bio-adhesive has a higher adhesiveness compared to existing treatments and burst pressure threshold approximately 6.4 times higher than normal intraocular pressure.
- In vivo rabbit trial showed that compared to cyanoacrylate glue, the bio-adhesive was safer, faster healing and led to less pain in rabbits.

## 1. Introduction

Tissue engineering has previously been employed within ophthalmology to generate implants or structures to replace partial or entire eye structures [1]. In recent years, 3D bioprinting has emerged. Bioprinting delivers biomaterials that can be aggregated to form a 3D structure based on computer-aided designs (reviewed in [2]). Biomaterial characteristics and the application surface will drive the printing process, for example, the cornea as a transparent tissue ideally requires a transparent biomaterial whilst retaining significant cell compatible and strength properties. Understanding the mechanical properties of the intended biomaterials therefore is essential for choosing, developing and fine-tuning the operating parameters of any bioprinter.

In addition to generating a standalone tissue, printing biomaterials directly onto injured sites represents a relatively new approach for tissue repair [3]. Using a hand-held 3D printer referred to as a biopen, Duchi et al., were able to deliver cells incorporated within GelMa bioink directly to chondral defects to show early cartilage regeneration [3]. This technique is referred to as in-situ bioprinting. We believe that in-situ bioprinting, the direct deposition of therapeutic biomaterials to corneal wounds, can be an effective treatment option for corneal injuries. Similar to 3D bioprinting, the success of in-situ bioprinting is reliant on the co-development of biomaterials and a compatible printing device. In particular, adhesive biomaterials also called bio-adhesive in this study, will have significant benefits in in-situ bioprinting applications.

Our group recently discovered that by combining human platelet lysate (hPL) with low concentrations of fibrinogen and thrombin, a transparent fibrin-based gel can be formed [4]. Using hPL to treat various ocular surface diseases such as dry eye syndrome has previously shown positive results [5]. Fibrin glue is a common tissue adhesive that has been in broad ophthalmic use for decades (reviewed in [6]). However, typical fibrin glue is non-transparent and contains high concentrations of fibrinogen (approximately 20mg/mL) making the component highly viscous and incompatible with functional visual acuity. Despite its wide use in surgery, the use of fibrin-based glue/bio-adhesives for corneal repair using the current composition is challenging.

Our earlier work demonstrated that the newly developed hPL and fibrin biomaterials possess suitable mechanical properties that are compatible with extrusion-based printing processes, adhesive to corneal tissue and support corneal epithelial cell growth [4]. To maximise transparency, the fibrinogen concentration was reduced to 2mg/mL, the physiological concentration in blood [4]. In this current study, we aim to generate an in-situ printing system capable of delivering the hPL based bio-adhesive directly to treat corneal surface and full thickness wounds. In particular, we will examine the rheology behaviour of the bio-adhesive including its cross-linking profile, adhesiveness and burst pressure. The generated mechanical data will be used to design an extrusion-based delivery device. As a further step to confirm the potential practical application of the system, a pilot rabbit trial was conducted to evaluate the novel bio-adhesive’s safety and effectiveness in sealing small corneal perforations.

## 2. Materials and Methods

### 2.1 Generation of hPL based fibrin Ink

The hPL-based fibrin bio-adhesive had two parts. Part A consisted of 20% hPL (v/v) and 4mg/mL human fibrinogen (Merck, USA) in DMEM/F12 (Life Technologies, USA). Part B was made of 10U/mL human thrombin (Merck) in DMEM/F12. Part A and B were stored separated in −80°C, and thawed in a 37°C water bath immediately prior to use. Mixing parts A and B in a 50:50 volume ratio resulted in the formation of a perforation-sealing hydrogel.

### 2.2 Measurement of rheological behaviour of the bio-adhesive

To develop a suitable extrusion-based printer, it remains important to perform rheological studies to measure the shear thinning properties of bio-adhesive, as well as to understand the cross-linking profile of the mixed material, which is the time required for gel formation after extrusion. Rheological behaviour was examined by an AR-G2 rheometer (TA instruments, USA) using a 40mm/2° cone plate geometry. Oscillatory tests were performed at 0.1% strain and 0.1 Hz frequency.

To measure the effect of shear, viscosity (η) of each part (Part A and Part B before mixing) was measured across a shear rate range of 0.1 to 500 1/s. Experiments were conducted at room temperature (RT) and 700 μL of each component was used during each experimental repeat. Minimum 3 repeats were performed.

Bio-adhesive crosslinking behaviour was measured at RT and 34 °C (to mimic corneal surface temperature) through an oscillatory time sweep test. Part A (500 μL) was initially added to the sample stage, followed by part B (500 μL). The experiment began immediately after the addition of part B to part A, and storage modulus (an indication of gelation) of the mixed sample was measured over 20 min. Minimum 3 repeats were performed.

### 2.3. Measurement of adhesiveness of the bioink

Porcine skin (Coles, 2246384P) was firstly scraped to remove the lipid phase and cut into pieces (2.0 cm × 1.0 cm), before soaking in PBS overnight at 4°C. Tissue samples were then glued onto plastic fixtures using cyanoacrylate glue. Part A (40 μL) was firstly applied to the skin, then 40 μL of Part B was added to part A. Finally, another piece of porcine skin was added with an overlapping area of ∼1.0 cm × 1.0 cm before the sample was incubated at 37 °C for 3 min. A lap shear test was conducted using the EZ-S mechanical tester (Shimadzu EZ-L) equipped with 10 N load cell and 1 mm min−1 crosshead speed (n ≥ 3) (Fig 1). The adhesive strength (E) was calculated using the equation E = F/S (where F is the failure stress and S is the bond area). Artiss glue (Baxter, USA) was used as a control following the same procedure. The test was repeated 3 times.

**Figure 1.**
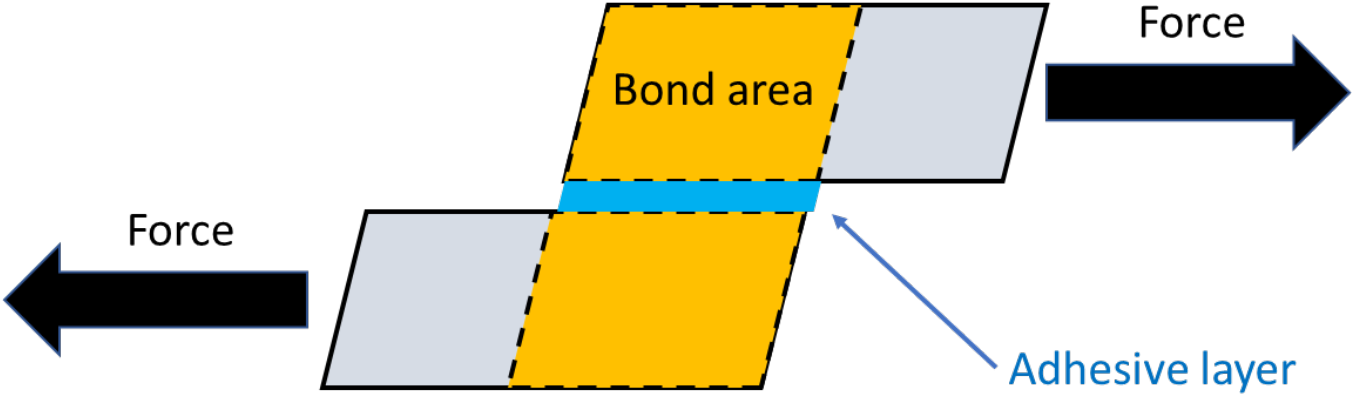
Schematic illustration of adhesion test.

Data were analyzed using ANOVA with Bonferroni multiple comparison post hoc test using SPSS Statistics (IBM, USA). P-values <0.05 were considered significant (*P≤0.05, **P≤0.01, ***P ≤ 0.001, n=3).

### 2.4 Evaluating burst pressure in an ex vivo human corneal model

Human corneas not meeting transplantation quality requirements were obtained from New South Wales Organ & Tissue Donation Service (Human Research Ethics Committee approval 14/275). A puncture wound of approximately 1.5mm in diameter was created centrally in the donor corneas. Corneas were then placed in a Hanna artificial anterior chamber (AAC). The pressure at which perforated corneas with no sealant exhibited stable leaking was measured for each cornea prior the test, and taken as the baseline pressure. The bio-adhesive (n=3) was administered to the wounds through the printing device for 5 sec and left for 5 min allowing adequate curing prior to burst pressure testing. The ACC was connected to a syringe coupled with a load cell device and actuated by a Shimadzu mechanical tester to record applied force within the fluid system (Fig 2). Sealant failure was determined when wound leakage was observed. Burst pressure was calculated as in Eq 2. and recorded in mmHg.

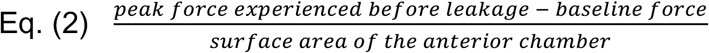

**Figure 2.**
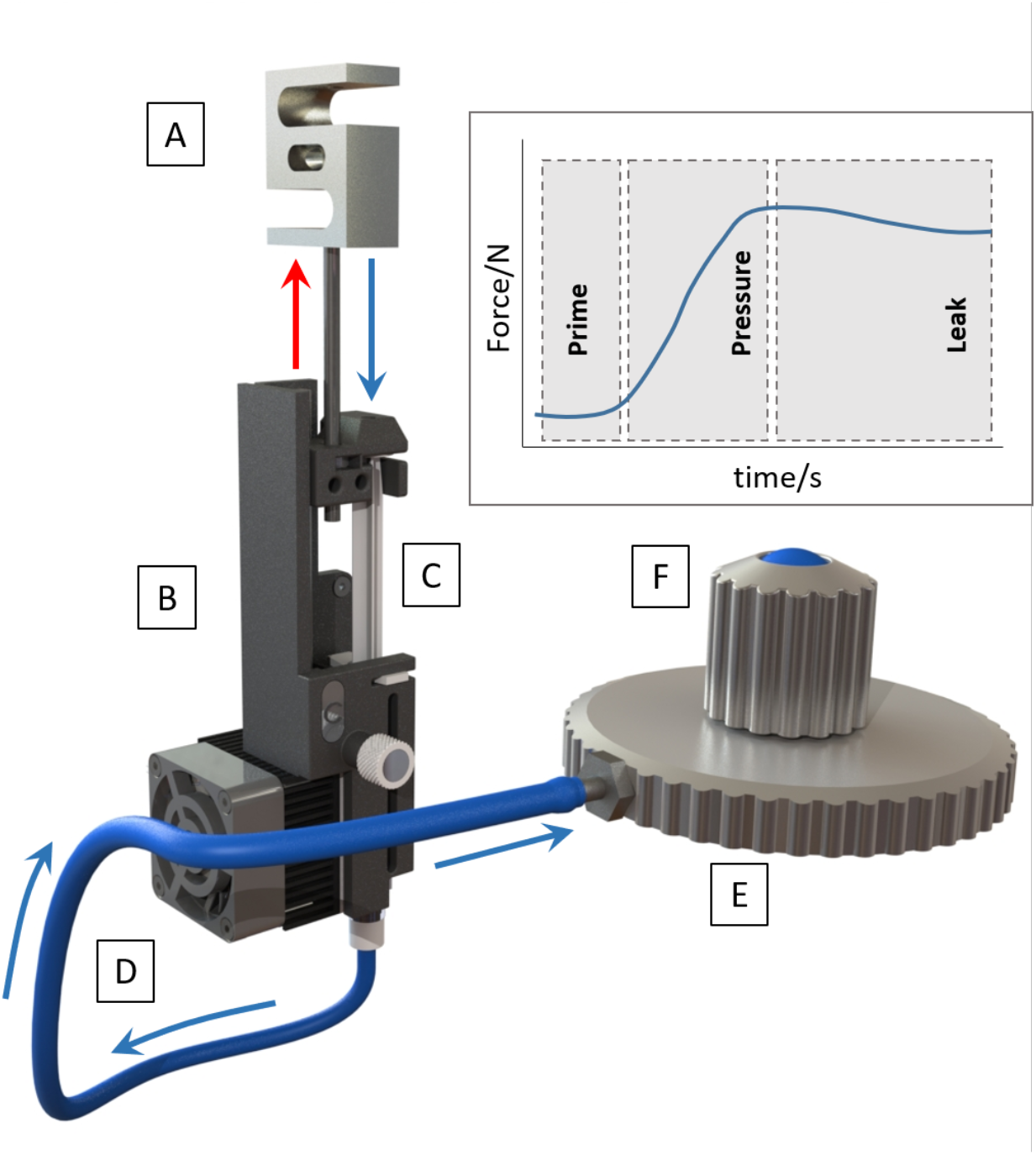
Burst pressure measurement system with inset indicating expected force profile observed during testing. A: Load cell coupled to Shimadzu mechanical tester. B: 3cc syringe mounting carriage rigidly mounted to test base of Shimadzu mechanical tester C: 3cc syringe containing water. D: Nylon tubing connecting syringe outlet to inlet of Hanna AAC. E: Hanna AAC. F: Cornea mounted in Hanna AAC.

### 2.5. Scanning electron microscopy of the hydrogel formed by the bio-adhesive

Scanning electron microscopy of the hPL and fibrin-based hydrogel was performed by using a JSM-6490LV scanning electron microscope (SEM) (JEOL Ltd., Japan). Scaffolds were frozen in liquid nitrogen for 40 seconds and then assessed directly. Images were taken at magnification (×200, ×500).

### 2.6. Development of hand-held biopen

An electromechanical two-component ink delivery device (Fig 3) was developed for controlled mixing and delivery of Part A and Part B ink components in a 50:50 ratio at a fixed target extrusion rate of 1μL/s. Extrusion force was applied to the plungers of the two separate 1 mL syringes via a geared micro-stepper motor (100:1 gear ratio) connected to a 4 mm lead screw with a thread pitch of 0.7 mm. Linear guides on either side of the plunging plate ensured consistent vertical motion. The motor was configured for both forward and reverse directions using a direction toggle switch, allowing for extrusion and subsequent resetting of the plunger position before next use. User control was achieved through depression of the micro switch located at the end of the pen. Continuous depression was required to drive the motor. The rate of motor revolution and in-turn translation of the plunger carriage was controlled by an Arduino Nano and A4988 stepper-motor controller after pressing of the pen activation push button.

**Figure 3.**
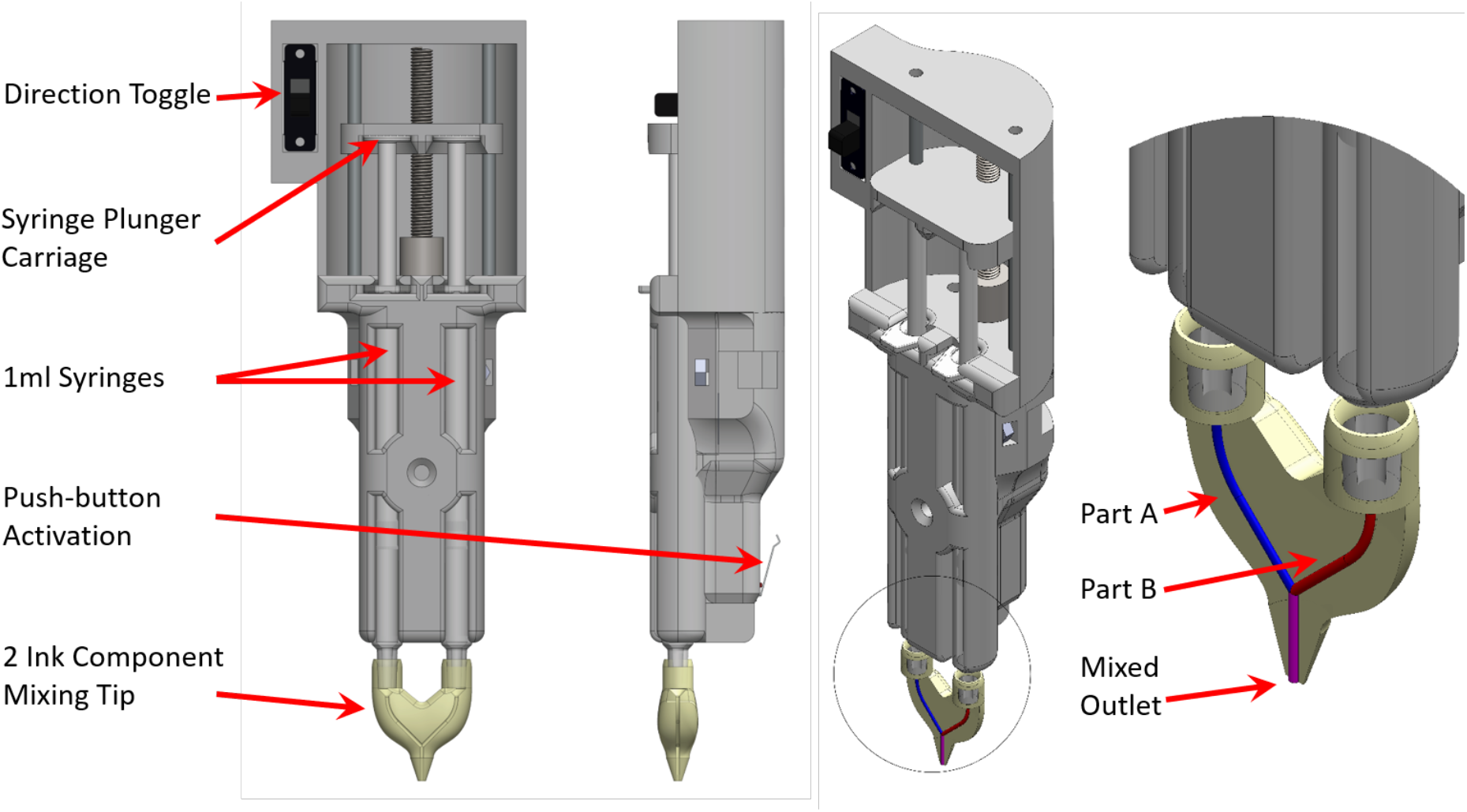
Schematic representation of electromechanical ink delivery device, iFix Pen.

### 2.7. *In-vivo* corneal perforation rabbit trial

Five female New Zealand white rabbits between 2 to 3kg were used in the trial. The trial was conducted at the University of Sydney Charles Perkins Centre Laboratory Animal House Facility with Animal Ethics Committee approval (AEC: 2018/1317). The procedure adhered to the ARVO Statement for the Use of Animals in Ophthalmic and Vision Research. Upon arrival, all rabbits were subject to a complete general health and ocular examination and were excluded if demonstrating signs of pre-existing corneal abnormalities. The rabbits were group housed and underwent acclimatisation for a minimum of three weeks prior to study commencement. Rabbits underwent the surgical corneal procedure under general anaesthesia performed by veterinary staff, followed by standard anaesthesia procedures.

The rabbits were divided into two groups to receive management of a surgically-induced corneal perforation with either bio-adhesive (treatment group), or cyanoacrylate glue (control group). The first group consists of three rabbits, two treated with bio-adhesive and one received the control glue treatment. The 2^nd^ group received modified surgical procedure (described as below) and was consisted of two rabbits, one for bio-adhesive treatment and one for control. In total, three animals received the bio-adhesive treatment and two received the control glue treatment.

#### 2.7.1 Perforation procedures

To create a full thickness perforation in the first group, the ocular surface and periocular skin surface region were prepared with 10% povidone-iodine solution, flushed with sterile saline (0.9%) and dried by sterile surgical spears. A sterile blade was used to create a complete perforation of approximately 2mm in length in the peripheral region of the cornea. The bio-adhesive was then extruded over the perforation in the treatment group (n = 2) and left to set for 2min. Commercial cyanoacrylate glue was administered over the perforation in the control group (n = 1). A modification of the procedure was made to the 2^nd^ group by dropping 10% povidone-iodine to 0.2% to minimise cytotoxicity. All wounds were created and treated as described previously. One control and one treatment in the 2^nd^ group.

#### 2.7.2 Post-surgery procedures

Following recovery from anaesthesia, rabbits were individually housed and received meloxicam 1.5 mg kg^−1^ subcutaneously daily. Topical chloromycetin eye drops and preservative-free eye lubrication were applied every four hours during working hours. Pain was scored every two hours by veterinary staff throughout working hours based on an in-house rabbit simple descriptive pain score system (Appendix -1). The scale was zero to four, with zero representing the best possible score. If a rabbit was scored 3 or higher, rescue analgesia was administered (buprenorphine 50 μg kg^−1^ IM or methadone 0.5 mg kg^−1^). Medication was continued until corneal wound healing was complete.

Corneal imaging (via portable slit lamp) following administration of topical fluorescein in the operated eye occurred four times per day at regular intervals until wounds healed. Complete wound healing was defined as the absence of fluorescein corneal staining following administration. The wound and surrounding area were examined for evidence of postoperative complications. Antibiotic eyedrops were administered following slit lamp examination.

#### 2.7.4 Histologic Examination

After wound healing, rabbits were monitored daily for side effects. At the three-week postoperative timepoint, rabbits were humanely euthanised with pentobarbitone IV in accordance with University of Sydney Laboratory Animal Services Protocol. Each rabbit was enucleated bilaterally and the eyes fixed with 4% PFA. Once fixed, corneas were dissected from the globe and then bisected and embedded in paraffin wax. These blocks were sectioned at a thickness of 5 μm and stained with hemotaxylineosin as previously published [7]. Histologic examination was performed with light microscopy.

## 3. Results

### 3.1 Rheology results of viscosity and cross-linking behaviour of a bio-adhesive

Both parts of the bio-adhesive showed shear thinning behaviour characteristic of non-Newtonian fluids (Fig 4). Viscosity became less dependent to shear rate due to gradual breakage of molecular networks under increased shear force.

**Figure 4.**
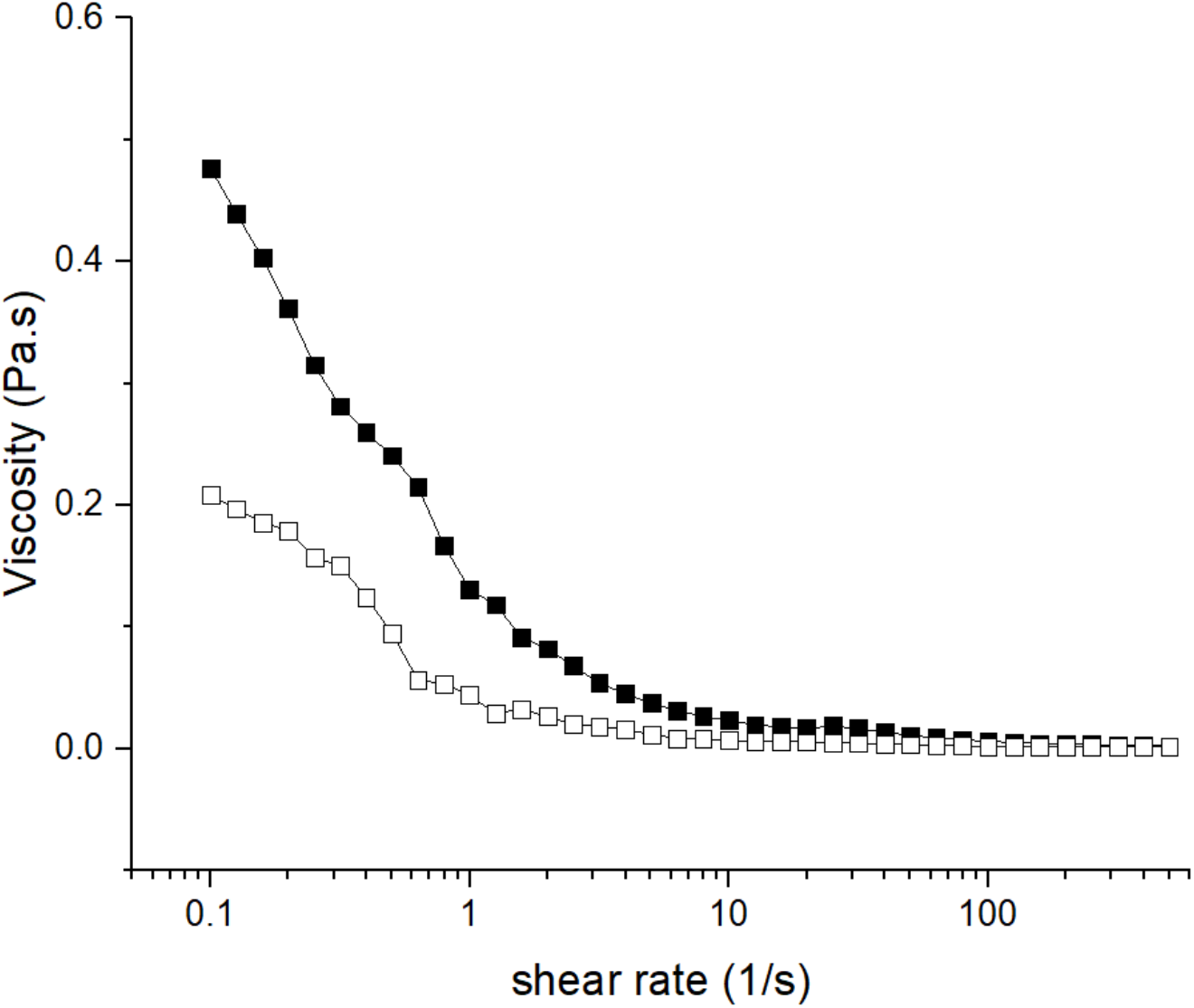
Variation of viscosity vs shear rate for part A (solid square) and B (hollow square).

Crosslinking behaviour of the ink versus time and temperatures was recorded 30 seconds after adding Part B to part A. Crosslinking was initiated as evidenced by an increase in storage modulus (G’) value, which gradually levels off by time indicating formation of crosslinked networks (Fig 5). When measured at 34°C, G’ increased dramatically after addition of part B, and crosslinking was almost completed after 2min30s. At room temperature, the G’ was still increasing after 2min30s until the end of experiment, albeit at a slow rate. The final G’ was higher at RT (8.7 Pa) than at 34°C (5.8 Pa).

**Figure 5.**
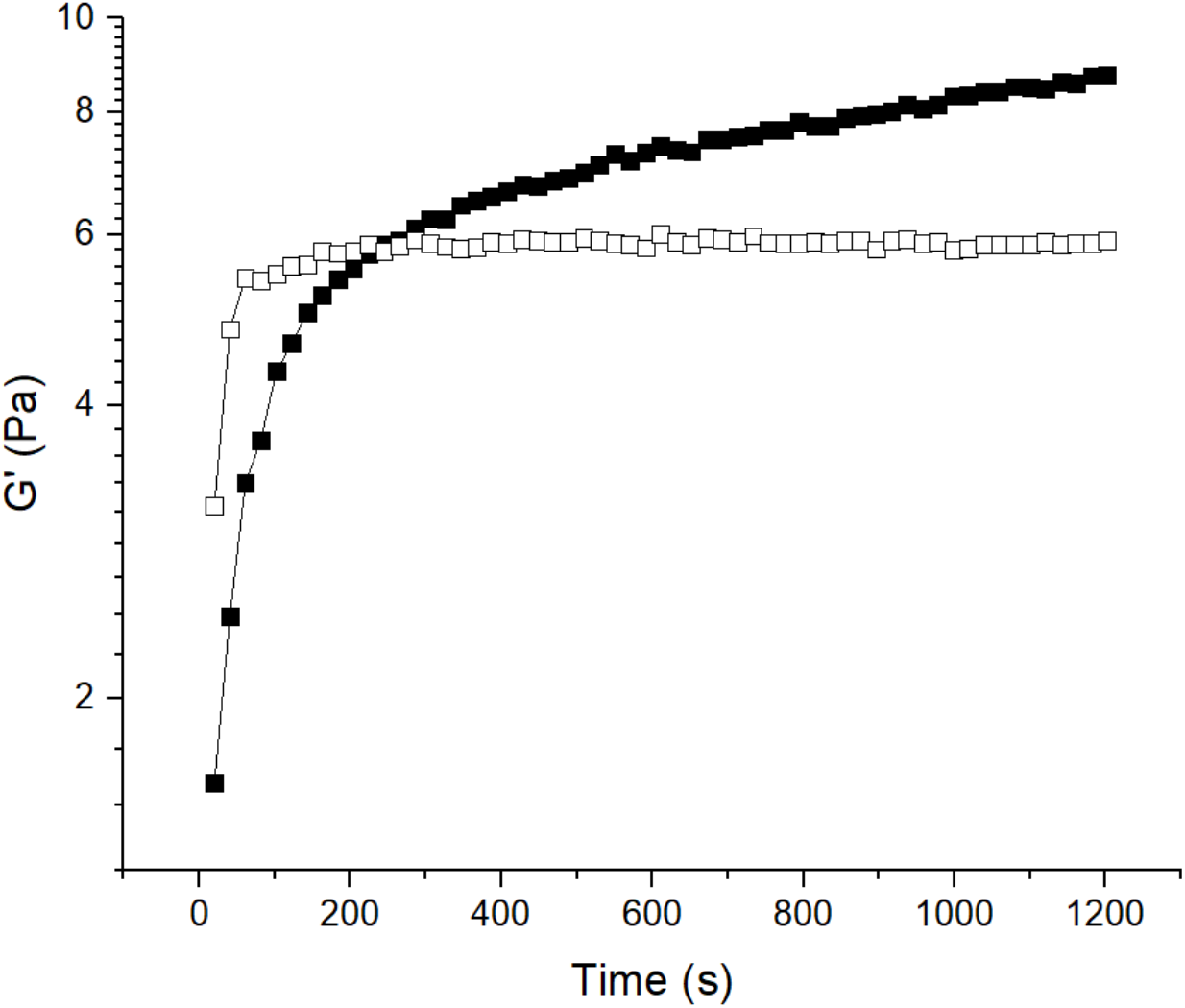
G’ versus time at RT (solid square) and 34 °C (hollow square).

### 3.2 Physical property results – adhesive and burst pressure

The bio-adhesive had an adhesive strength at 1575 ± 195 Pa higher thant Artiss^®^ (control) at 1260 ± 228 Pa, despite not being significantly different. Mean burst pressure of the bio-adhesive to seal a 1.5mm perforation was 91 ± 27mmHg in the human corneas. This pressure exceeded the mean intraocular pressure of a normal adult eye (approximately 14mmHg) [8–10].

### 3.4 Microporous structure of the hydrogel

Throughout the cross-sectional area, hydrogel formed by the hPL based fibrin bio-adhesive exhibited highly porous, micro-structured interconnected networks that were formed by fibres (Fig 6). SEM images showed multiple fibrin fibres assembled and bundled together. The diameter of each fiber ranged from ∼0.1 to 0.5 μm, consistent with diameters of previously reported fibrin fibers [11].

**Figure 6.**
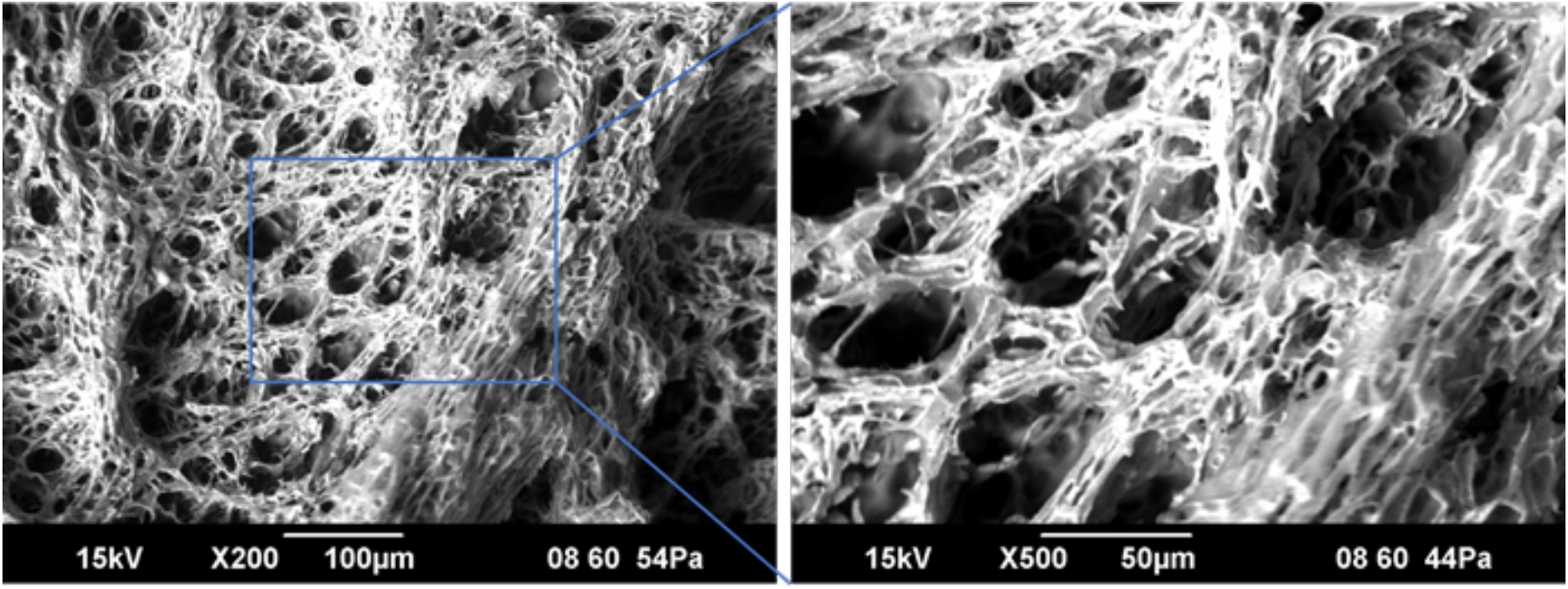
SEM photographs of hydrogel structure formed by the bio-adhesive.

### 3.5 Extrusion based electromechanical pen for bio-adhesive delivery

Flow rate testing was conducted to determine total output of the delivery device. One syringe filled with water was driven for 30 seconds, and the deposited volume of water measured. Measured flow rate over six trial replicates was 1.02 ± 0.03 μL/s and effective within a 5% tolerance of the target volumetric output. This trial was then performed using the two component inks with a target flow rate of 0.5 μL/s for each ink. Resultant mean extrusion rate was 0.512 μL/s with a deviation < ± 0.01 μL/s showing very small discrepancy.

### 3.6 In-vivo test results

The two bio-adhesive treated rabbits in the 1^st^ group showed moderate conjunctival inflammation and developed secondary corneal ulceration (Fig 7A). The control rabbit treated with glue developed severe conjunctivitis, chemosis and secondary corneal ulceration. Secondary corneal ulcers healed more quickly in the bio-adhesive treated rabbits, and the corneal perforation healed approximately 3 times faster in the bio-adhesive treated cornea (Fig 7A).

**Figure 7.**
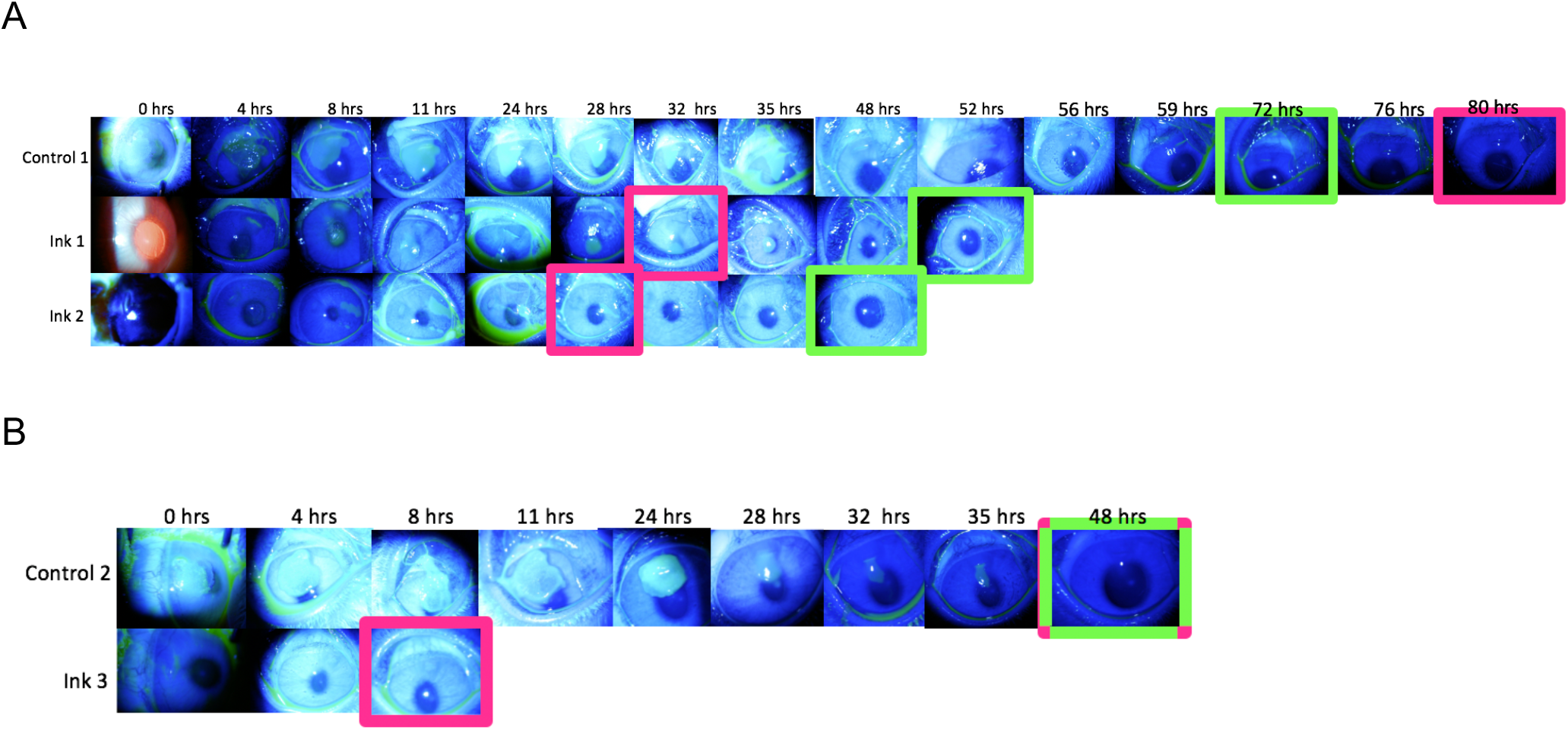
Two rounds of corneal perforation tests. A: 1^st^ round of test showed that bio-adhesive treated groups healed much quicker than control cornea. B: Ink treated cornea had no complications and healed 6 times faster than control. Pink rectangle box: perforation. Green rectangle box: ulceration.

Cyanoacrylate glue is toxic to epithelial cells, therefore development of secondary corneal ulceration could be an expected complication in the control rabbit. However, it was an unexpected complication that secondary ulcers would develop in the bio-adhesive treated corneas given the nature of the bio-adhesive. Reduction of povidone iodine (0.2% vs original 10%) was made to reduce potential cytotoxic when preparing the ocular surface for surgery. With the modification, the bio-adhesive treated cornea showed no macroscopic evidence of inflammation or complications, whereas the control cornea still developed 2^nd^ corneal ulceration (Fig 7B). The corneal perforation treated with bio-adhesive healed at 8 hours postoperatively, six times faster than the control cornea (Fig 7B).

In the first group, bio-adhesive treated rabbits had lower pain score and received less rescue analgesia in comparison to the control rabbit (Fig 8A). In the 2^nd^ group, no additional analgesia was required for the bio-adhesive treated rabbit, while the control rabbit required an additional four doses of rescue analgesia postoperatively (Fig 8B).

**Fig 8:**
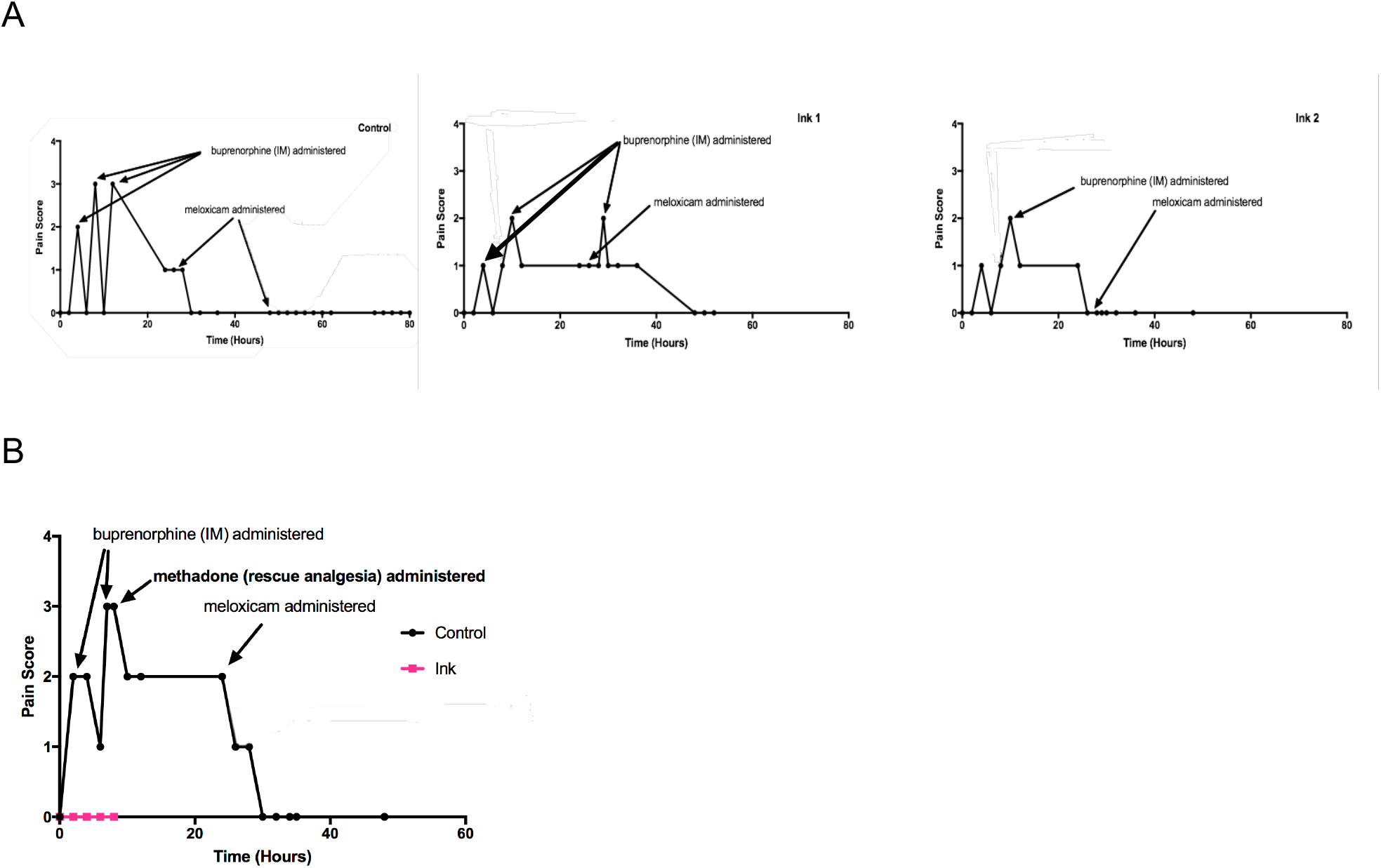
Pain score and analgesia administration. A: First round of rabbits. B: Second round of rabbits.

The H&E microscopic analysis showed similar tissue structures across all corneas harvested 3 weeks after surgeries. Both bio-adhesive and control corneas showed regenerated epithelial layer and stromal scar formation shown as a dense stromal structure compared to the surrounding normal tissue (Fig 9).

**Figure 9.**
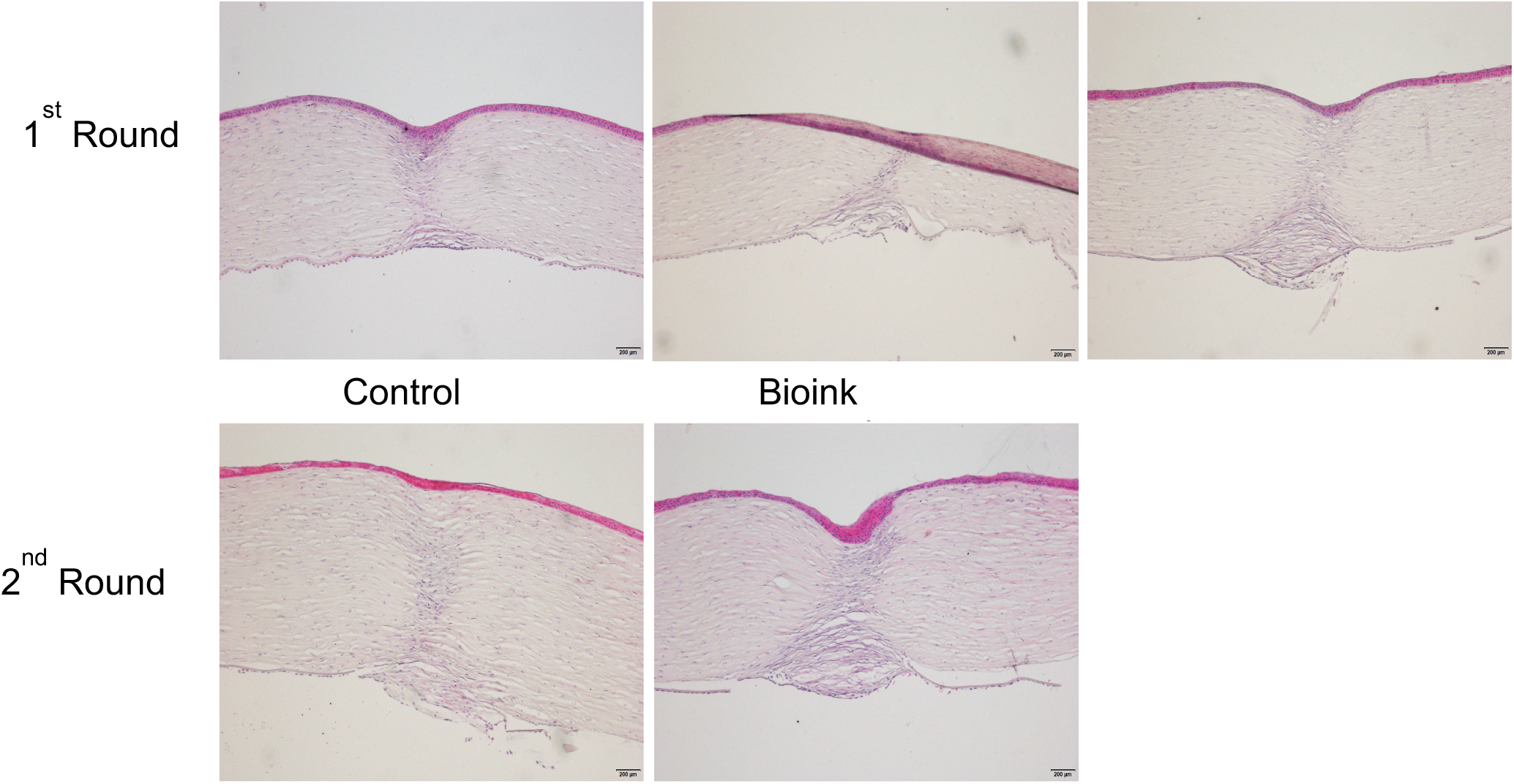
H&E images of tissue structures of perforation region.

## 4. Discussion

In this study, we successfully developed a novel, *in-situ* bio-adhesive-based printing system to treat small corneal perforations. Our bio-adhesive was effective, indicating a 6x faster healing rate and zero pain score when compared to control (cyanoacrylate glue). The bio-adhesive used was an hPL-based fibrin biomaterial previously developed by our group which showed high transparency and cell compatibility [4].

The crosslinking profile of the bio-adhesive showed that gelation started immediately after mixing, and full cross-linking achieved around 2min at 34 °C (the surface temperature of cornea). The microstructure of the hydrogel formed showed an organised porous structure formed by fiber bundles, which was different to published fibrin gel structures that have found single fibrin fibers interconnected in a mesh format [11]. We also compared its adhesiveness to the Artiss glue that is currently used in clinics and showed that our bio-adhesive had higher average of adhesiveness strength, albeit the difference did not reach statistical significance. In corneal treatments, Artiss glue is frequently used to seal small perforations therefore this outcome indicated potential that our bio-adhesive may be used in similar application.

A hand-held extrusion device was designed based on the mechanical properties of the bio-adhesive to have an extrusion rate of 1μL/s. This extrusion rate was indicated to provide sufficient operation time for surgeons to apply the ink to the wounded site whilst minimising the risk of the bio-adhesive clotting in the nozzle. In this trial, the novel hand-held printing device successfully delivered bio-adhesive to the injured sites. The bio-adhesive solidified in 2 minutes, remained on the cornea even after instillation of antibiotic eye drops with no leaking identified.

Corneal perforation is associated with severe pain, infection and may lead to blindness [12–14]. Management depends on the size of the perforation. As a first trial of this novel system, the aim was to demonstrate the usability of the bio-adhesive and delivery device by surgeons and to evaluate the safety and efficacy of the novel bio-adhesive compared to standard clinical care. The size of the perforation, remains a relative limitation of the study however despite the small perforation, in all rabbits, the bio-adhesive treated corneal surface healed more rapidly compared to controls treated with cyanoacrylate glue; and all bio-adhesive treated rabbits had lower pain scores than the control rabbits. In the last two rabbits, the bio-adhesive treated cornea showed imminent advantages to cyanoacrylate glue with the wound completely healed 8 hours postoperatively, six times faster than control. This rabbit had a consistent pain score of 0 with no additional rescue analgesia required. This trial suggested that the bio-adhesive could lead to faster healing, alleviate pain caused by corneal injuries either directly or indirectly, by avoiding the conjunctival inflammation induced by cyanoacrylate glue. Histological data showed that in all cases, scar structures can be seen in the injured area with no obvious morphology differences observed between the bio-adhesive treated and control group. All corneas were harvested three weeks after surgery and no complications were observed in the chronic phase of the study once the corneal perforations had healed suggesting longer-term potential. Although promising, our rabbit trial had some limitations. The rabbit trials were conducted over two different periods due to the unexpected corneal ulceration in the bio-adhesive treated group in the initial rabbits. The 2^nd^ corneal ulceration in the bio-adhesive treated rabbit was resolved by using less concentrated 0.2 % povidone-iodine. The reduction was suggested by the veterinarian due to the lack of a Bowman’s membrane in rabbit’s cornea which have made the corneal structure more susceptive to insult of povidone-iodine than human cornea. However, the sample size was small. As discussed previously, the perforation created was a linear 2mm wound which can be self-sealed. It is worth noting that the burst pressure test was performed successfully on a 1.5 mm circular wound that was not self-sealing. Further in vitro studies with more significant, repeatable wound sizes and a larger sample size would be more appropriately for evaluating the sealing effectiveness of this bio-adhesive.

A number of novel treatments have been published as alternatives to treat corneal perforations including using collagen-based fillers [15], fibrin glue-assisted amniotic membrane [16], and LiQD cornea [17]. These studies have demonstrated successful sealing results. LiQD cornea also contained fibrinogen and thrombin, similar to the fibrin glue-assisted amniotic membrane approach to allow for in-situ gelation. Both collagen-based filler and LiQD cornea required additional steps including the addition of LP-PEG, and DMTMM to aid gel formation. Increased temperature to 50°C for mixing DMTMM was required in LiQD method. Our approach required no additional chemical crosslinker and represents an in-situ printing system, which is a completely different approach. The extrusion-based device allows in-situ gelation as per the LiQD treatment, but also allows in-situ printing with no implant or membrane required. This relatively simple model increases the practical potential of the device, in particular in developing countries that may not have access to trained medical personnel. In addition, both studies using CLP-PEG and LiQD did not report pain assessment. We have identified obvious pain reduction in rabbits treated with our bio-adhesive. Reducing pain and discomfort following treatment in the human patient would represent a potential significant advantage. Furthermore, as no additional chemical cross-linking was used and no requirement of high temperature, our bio-adhesive has potential to incorporate bioactive molecules such as growth factors, antibiotics and even cells. Further studies on this direction will be conducted.

In conclusion, this study suggested that the bio-adhesive was safer than cyanoacrylate glue to treat small cornea perforations. Despite the similar scar structures formed, the bio-adhesive resulted in faster recovery and considerable pain reduction. However, this study has limitations. The 2mm wounds created were a slit wound, therefore there was the possibility of the tissue self-sealing postoperatively. A trephined circular wound should be examined. Furthermore, bio-adhesive alone may not be able to address a larger perforation due to the minimum 2 minutes gelation time. Nevertheless, we can report a novel device and approach to treat small corneal perforation with potential to accelerate healing and addressing pain without additional management. The bio-adhesive composition was simple with only three active ingredients which are human-derived and clinically-safe reagents. The delivery system is a hand-held device, which is easy to deliver and store. This combined device and bio-adhesive represents a potential simple, effective alternative to cyanoacrylate and fibrin glue for small corneal perforation treatment.

## 5. Acknowledgement

We thank Australian National Fabrication Facility (ANFF Materials Node), Translational Research Initiative for Cell Engineering and Printing, Australian Research Council (ARC) Centre of Excellence Scheme for the use of facilities at the University of Wollongong Electron Microscopy Centre (EMC facilities), and the support of the ARC Industrial Transformation Training Centre in Additive Biomanufacturing (IC160100026). We acknowledge the funding support from Australian Research Council (ARC) Centre of Excellence Scheme (CE140100012), Sydney Eye Hospital Foundation and NSW Medical Device Fund. The hPL used within the study was a gift from Australia Red Cross. We also thank Mr Jordan de Barros for the development of the mechanical pressure test rig used in this paper.

